# Astrocytes control UP state and slow oscillation periodicity in human cortical networks

**DOI:** 10.1101/2024.11.01.621156

**Authors:** James A Crowe, Amjad H Bazzari, David A Nagel, Sergei G Sokolovski, Edik U Rafailov, Eric J Hill, Rhein H Parri

## Abstract

Astrocytes are well known for homeostatic roles in energy maintenance, neurotransmitter recovery and immunoreactivity, however their contributing role in active modulation of neuronal activity remains controversial. Recent evidence highlights astrocytes as a potential signalling partner in ongoing high-order neuronal activity, and a modulator of baseline activity.

Here we utilise an iPSC-derived in vitro cortical network model to describe a slow-wave oscillation and examine the contributions of astrocytes to development of this oscillatory activity, dependant upon generation of UP/DOWN state phenomena. To examine the role of astrocytes we invoke an acetylcholinergic oscillation in the neuronal population by addition of carbachol, which proved to be a robust mechanism for eliciting prolonged and synchronised network bursting. Pharmalogical interrogation determined oscillatory maintenance was dependent on both glutamatergic and purinergic signalling pathways. Upon further interrogation, we determined that astrocytic calcium signalling was essential to timing of the oscillatory signal. By utilising chemogenetic actuators we showed that whilst neurons are essential and sufficient for instigating the oscillatory signal, astrocytes played a key role in timing.

This previously unreported mechanism may contribute to initial development of brain activity, and may underlie a basal activity present later during adulthood. In collaboration with recent results highlighting the role of active gliotransmission, this highlights astrocytes as an important research target for understanding brain activity alterations during development and disease.

## Introduction

Oscillations within neuronal networks are a commonly investigated feature of mammalian brains, often associated with varying states of vigilance, consciousness and disease ^1,2^. Monitoring of *in vivo* human brain activity is primarily achieved via analysis of these oscillatory waves using non-invasive techniques, such as electroencephelogram (EEG), which are routinely conducted both experimentally and clinically, to investigate brain states such as sleep, awareness, and seizurogenic onset. In pathological states such as Parkinson’s disease, Alzheimer’s disease, epilepsy, and neonatal hypoxia, exhibition of characteristically altered oscillatory frequencies can highlight major network dysfunction, e.g. often a loss of more complex activity to a prominent oscillatory ‘tone’ ^3–9^). However, clues to the cellular mechanisms underlying manifestation of these oscillations have, by necessity, been largely obtained from experiments using *in vivo* and *ex vivo* anatomically intact rodent models ^10,11^. Despite rodent models exhibiting many phenomenological features of human brain activity, it is unclear to what extent mechanistic data from rodent tissue can be extrapolated to humans due to the higher complexity of individual cell types (e.g. astrocytes) and their functional differences when comparing human versus rodent neocortical tissues ^12,13^.

To decipher the cellular and synaptic mechanisms that underlie human brain oscillations, we therefore require robust, interrogatable, human models that scale from individual cells to globally-active networks. Whilst long established that human iPSC-derived (hiPSC) neurons display appropriate protein expression patterns and exhibit comparable single neuron electrophysiological properties to *ex vivo* tissue ^14–17^, it remains scientifically important to determine to what extent hiPSC-derived neuronal networks can generate sophisticated and complex physiological activity. This key feature defines their ultimate utility as functional models of *in vivo* human brain networks for understanding neurodevelopment and disorders of neuronal function.

Encouragingly, recent advances using hPSC-derived cortical organoids implicate that complex 3D tissue-like cortical networks can exhibit network-level oscillations in local field potentials ^18–20^ resembling activity from intact adult brain preparations and foetal brain development, including disease-replicative epileptiform activity. A selective advantage of using hPSC-derived cell models over rodent *ex vivo* tissue is that neural cell subtypes can be selectively differentiated and investigated in isolation, or in combinations with other cell types, enabling cell-, synaptic- and network-specific physiological interrogation. This has been demonstrated by transcription factor-induced cell modelling, and interconnecting brain-regionalised assembloids, and defined generation of region-specific neuronal subtypes shown to be therapeutically relevant in novel treatment schemes for neurodegenerative disorders ^21–24^. Yet despite these advances little remains known about the specific cellular and synaptic mechanisms underlying the generation of such activity phenomena.

To determine whether hiPSC-derived network models are capable of mechanistically interrogating higher order activity, we must focus our initial search on established, fundamental oscillatory phenomenon of the mammalian brain, such as the generation of slow-wave oscillations. “Slow” oscillations exhibit periodic membrane potential changes at a frequency of < 1Hz and are observed primarily *in vivo* during NREM sleep states, anaesthesia, and reported in “default” network activity in isolated cortical tissue ^25,26^. Prior animal model experiments characterise slow oscillations by distinctive UP and DOWN states in neuronal membrane potential, and report a common initiation site in neocortical layer 5 before propagation to other cortical layers ^26–29^. Remarkably they are identified as early as the onset of electrical activity during neurodevelopment, in both human and rodent *ex vivo* cortical tissue ^30^ implying a key role in early brain development. This is evidenced by the development of sporadic synchronised infra-slow *tracé discontinu* (GW22-30) activity during *in vivo* prenatal EEG studies, which eventual becomes more consistent over time with *tracé continu* and then further complexifies into *tracé alternant* (GW34-44), before settling into traditional slow wave sleep and spindles. In animal models this has been reported by the development of cENOs, GDPs and SATs ^31^. Importantly, hypoxic injury can decrease the complexity of prenatal brain activity resulting in burst-supression EGG patterns (BSP).

We determined to mechanistically interrogate oscillations in an *in vitro* human cortical model combining hiPSC-derived lower layer 5/6 cortical neurons and astrocytes ^15,32^. Using a comparative approach of live imaging (calcium indicators, glutamate reporters), electrophysiological recording (patch clamp, MEAs), chemogenetic actuators (Gq-DREADDs), and defined pharmacological intervention, we report that human cortical layer 5/6 networks respond to muscarinic cholinergic receptor activation by exhibiting highly periodic network oscillations.

Observed individual cell unit activity and overall network oscillation recapitulated many key features of UP and DOWN states, and characteristics of slow wave oscillation reported in rodent models and human tissue ^26,28,33^. Our data indicated a dependence on adenosine A_1_ receptor activation, and importantly, astrocytic calcium signalling for oscillation periodicity ^34,35^. Despite astrocytic control over periodicity, neurons remained the essential driving force of the oscillatory activity. Our results establish hiPSC-derived networks maintain the capacity of human cortex to exhibit fundamental types of oscillatory activity that depend on neuron-astrocyte interactions, and that highly-purified subtype cultures are amenable to mechanistic interrogation. These data suggest a novel role of neocortical astrocytes as active maintainers of slow wave oscillatory activity, previously implied by their role in maintenance of circadian rhythm, and development of visual cortex, and adding to recent reports of active gliotransmission that modulates neuronal activity ^36–40^. Importantly, this study also identifies a “missing link” in many computational neuronal models, an astrocytes “clock” to aide in timing neuronal oscillations alongside conventional interneuron components.

## Methods

### Cell culture

Human iPSC-derived cortical neural progenitor cells (NPCs) were obtained from Axol bioscience (Newborn, 0 years, Male, ‘Healthy’ donor). Thawed cells were resuspended in 200 μL cm^−2^ Neural maintenance media (Axol Bioscience) supplemented with 10 μM Y-27632 dihydrochloride (Selleck Chemicals), and then plated at a density from 100,000 to 150,000 cm^−2^ in 6-well tissue culture plates pre-coated with SureBond (Axol Bioscience), or Poly-L-ornithine (Sigma-Aldrich) + 20 ng mL^−1^ murine Laminin (Sigma-Aldrich). Cultures were then incubated in a 37 °C, 5% CO_2_, 95% atmospheric incubator overnight. After 24 hours (hrs) the media was replaced with 200 μL cm^−2^ fresh Neural maintenance media, without Y-27632 dihydrochloride. Media was replenished every 48 hrs until passaging when cells reached 90% confluency. 48 hrs prior to passaging 10 ng mL^−1^ recombinant human FGF2 (Axol Bioscience) was added to promote proliferation, with total FGF2 treatment per batch limited to < 6 days to prevent off-fate differentiation. SureBond coating consisted of 1:50 dilution of SureBond in DPBS without Ca^2+^ or Mg^2+^ (Sigma-Aldrich), incubated for 24 hrs at 37 °C. Poly-L-ornithine coating was diluted 1 mL in 4 mL distilled H_2_O (Sigma-Aldrich; dH_2_O), coated wells at 200 μL cm^−2^ and incubated for 2 hrs. After incubation Poly-L-ornithine was rinsed twice with dH_2_O and incubated with 200 μL cm^−2^ laminin diluted in dH_2_O and incubated for 24 hrs prior to use.

To passage neural progenitors, media was removed, and cultures were briefly rinsed with DPBS without Ca^2+^ or Mg^2+^ (Sigma-Aldrich) before incubation with StemPro Accutase (Gibco) for 5 mins at 37 °C. The culture was then gently pipetted until fully detached from the culture platform, and gently dissociated before accutase was neutralised with 2 x volumes of Neural maintenance media. Cell suspension was centrifuged at 200 x g for 5 mins to form a cell pellet. Supernatant was discarded before resuspension in 1 mL Neural maintenance media supplemented with 10 μM Y-27632 dihydrochloride. A cell count was performed and cells were re-plated on pre-coated plates at a density of 100,000 to 150,000 per cm^2^.

Human iPSC-derived Astrocyte progenitor cells (Axol Bioscience) were plated on to tissue culture plastic coated with SureBond-XF (Axol Biosciences) coating prior to culture. SureBond-XF was diluted 1:200 in DPBS without Ca2+ or Mg2+ and cultureware incubated for 4 hrs at 37 °C. Frozen cryovials were removed directly from cryogenic storage to a 37 °C waterbath for 1-2 mins until thawed. Cells were resuspended in pre-warmed astrocyte maturation medium and the cell suspension was centrifuged at 400 x g for 5 mins. After removing supernatant, the cell pellet was resuspended in pre-warmed Astrocyte maturation medium (Axol Biosciences) and were plated at a density of 25,000 cells cm-2 in 6-well tissue culture plates pre-coated with SureBond-XF. After 48 hrs fresh pre-warmed Astrocyte maturation media was fully exchanged every 2 days. For progenitor astrocytes, cultures were regularly passaged upon reaching < 90 % confluency for 2 weeks before use.

Post-expansion, whilst the progenitors are still mitotically active, cells were passaged and terminally plated onto SureBond + Readyset (Axol Bioscience), or Polyethylenimine (PEI) (Sigma-Aldrich) and murine laminin (Sigma-Aldrich) pre-coated 13mm ø, thickness 1 coverslips (VWR) at a density of 100,000 cm-2 in a 24-well plate for immunocytochemistry, calcium imaging and patch clamping.

For synchronous differentiation of NPC cultures the medium was initially exchanged with Neural maintenance media for a 24-48 hr period to allow normalisation of cells. Once at a sufficient density upon final plating, media was exchanged to BrainPhys™ + Neurocult™ SM1 (Stem Cell Technologies) supplemented with 10 ng mL-1 human recombinant BDNF (Stem Cell Technologies), 10 ng mL-1 human recombinant GDNF (Stem Cell Technologies) and 2 μM γ-Secretase Inhibitor XXI, Compound E (Calbiochem). Cultures were fed with freshly prepared BrainPhys™ + SM1 + BDNF + GNDF + Compound E every 72 hrs for 6 days until cells show a long neurite morphology and decreased proliferation. After completion of synchronisation on day 7 of Compound E addition, media was half-exchanged with BrainPhys™ + SM1 + BDNF + GDNF every 72 hrs until experimentation. For synchronised co-cultures, the synchronisation protocol was followed until day 11 post the initial addition of Compound E. On Day 12 post initial synchronisation with Compound E, astrocytes (Axol Biosciences) were passaged and resuspended into BrainPhys™ + SM1 + BDNF + GDNF and plated in a half media exchange at a density of 10,000 – 15,000 cells cm-2 and allowed to settle for 48 hrs. After this time cultures were exchanged every 72 hrs with half media changes of BrainPhys™ + SM1 alone until experimentation.

### Calcium imaging

Unless otherwise stated calcium imaging was always performed during a period from day 45 *in vitro* until a maximum of day 80 *in vitro*. Spent media was removed from cultures and they were loaded with 10 μM Fluo-4 AM (Invitrogen) in fresh BrainPhys™ media for 45 mins at 37 °C in an incubator (Gee *et al.* 2000). BrainPhys™ media was chosen due to its constitutive relevance to aCSF. Following incubation, cultures were rinsed for 5 mins in pre-warmed BrainPhys™ media before coverslips were transferred to a microscope setup for immediate imaging.

Carbenoxgenated aCSF (126 NaCl, 2.5 KCl, 26 NaHCO3, 1.25 KH2PO4, 1 MgSO4, 2 CaCl2 and 10 glucose (all values in mM, Fisher Scientific) was perfused via a peristatic pump (SciQ 400, Watson Marlow) onto the coverslip immediately after securing to the tissue perfusion chamber (RC-25, Warner instruments) with a silicon paste. Perfusion was maintained at least 5 mins prior to exposure for imaging. The perfusion bath was placed upon the stage of a Nikon FN1 microscope (Nikon). Perfusates were passed through a heat block set to 37 °C to ensure physiological temperatures. Upon acquisition MetaMorph imaging software (Cairn Research) was used to control initiation of a LED light source (OptoLED power supply) and an ORCA-05G camera (Hamamatsu). The software allowed initiation of 470 nm (49011, FITC filtered, Chroma) or 590 nm (ET555/25x ET632/60m, Chroma) excitation light sources via a 20x objective (Fluor, 0.50W, DIC, WD2.0, Nikon) with a timing dependent on entered values. Typical exposure was between 20 – 40 ms depending on the relative level of Fluo-4 loading, and imaging frequency was 1 frame every 2 seconds (0.5 Hz). A baseline recording prior to exposure to experimental conditions was typically acquired for 6 mins to ensure cultures were healthy and responsive. During experimentation, a further baseline would be taken for a given condition e.g. addition of carbachol to induce an oscillation. Cultures were treated with the compounds for pharmacological interrogation. Furthermore, a variant of the ACSF described was produced which was osmotically balanced by reducing to 123.5 mM NaCl but contained 5 mM KCl instead of 2.5 mM KCl, this is known hereafter as 5 mM KCl ACSF. Post-acquisition data was exported as a .tif metastack for further processing and analysis.

Metastack files from acquisition were imported into FIJI for further analysis. Initially a frame-shift correction was conducted to align the image across all recording periods and thus to ensure that regions of interest (ROIs) are spatiotemporally stable. This was achieved using the template matching plugin for FIJI (Tseng *et al.* 2011). A maximum intensity projection was produced by using the Z project function of FIJI on a metastack. This maximum intensity stack was then used to add ROIs for measuring mean pixel intensity. Upon generating ROIs the values of the mean pixel intensity across the z-stack were extracted and entered into Excel 2013 (Microsoft). Values were normalised to baseline using a ΔF/F0 * 100 calculation where ΔF represented current mean pixel intensity (FX) minus the lowest value throughout the time sequence (F0). Upon normalisation the percentage ΔF/F data were exported as CSVs for analysis in Clampfit 11.0.3.03 (Molecular Devices). In Clampfit, the baseline was manually adjusted via the Adjust > Baseline function to allow clear detection of fluorescent peaks from the original baseline. Once baseline corrected traces individually underwent manual peak detection and rejection via the Event Detection > Threshold Search function. The baseline was set to 0 % DF/F and the minimum threshold was set to 2 % DF/F to avoid detection of background noise. Once complete the detected events per trace were extracted to Excel 2013 for further handling. Excel (Microsoft office 2013) was then used to generate means and standard error of the mean for key outcomes e.g. frequency, inter-event-intervals (IEIs) and peak amplitude for individual events. Excel was also used to compute cumulative probability of pooled frequencies across samples.

### GECI / iGluSnFR imaging

Co-cultures and astrocyte monocultures were incubated in virus diluted in 300 μL media per individual 24 well (BrainPhysTM + SM1 for both neuronal and astrocytic conditions. Cultures were then incubated in their original maintenance media and exchanged every 2-3 days as required until 7 days post treatment. For co-cultures, cells were incubated in a 1:1 solution of conditioned media (previously collected, no viral load) and fresh media to ensure typical half-media exchange. After 7-14 days post treatment cells demonstrated sufficient expression of the constructs for imaging or experimentation.

### Multi electrode arrays

Co-culture of neurons and astrocytes on MED64 multi-electrode arrays was adapted from Odawara et al 2016. Carbon-nanotube electrode multi-electrode array probes (MED-R515A, MED64) were initially sterilised by submergence in 70 % ethanol in a 10 cm petri dish for 15 mins followed by three rinses in dH2O, and then air drying under a UV light source for sterilisation. A 0.05% Polyethyleneimine solution was used to pre-coat the MEA probe for 45 minutes at 37°C in an 5% CO_2_ / 95% O_2_ atmospheric incubator. PEI was then aspirated and rinsed three times with sterile distilled H_2_O. Coating was then performed with iMatrix-511 recombinant human laminin. Setup of MEA cultures was similar in process to terminal plating of cells for synchronised co-cultures. However, whilst MEAs were pre-coated with Readyset+SureBond/PEI+laminin cells were seeded in a different manner. Once the cells were passaged and in suspension, coating solution was removed from the probe well and a sterile glass cloning ring was gently placed at the centre of the probe covering the electrodes. Cell suspension was left at a high concentration, and cells were gently pipetted at 100,000 cells 100 μL-1 into the cloning ring. Immediately after, prewarmed neural maintenance media and 10 μM Y-27632 dihydrochloride was added into the well, making certain to add gently as to not disturb the cellular suspension in the cloning ring. The MEA was then gently transferred in their 10 cm petri dish to an incubator and left for 1 hr for the cells to settle. After 1 hr the cloning ring was removed gently and the MEA examined to ensure the cells were at 100 % density covering the electrodes. Cultures were left 24 hrs and then media was fully exchanged directly to BrainPhys™ + SM1 + BDNF + GDNF + Compound E. Hereafter, the protocol for co-culture was performed exactly as described above.

### Recording culture electrical field activity using a multi-electrode array

Once sufficiently matured MEA probes were sealed using a lid that allowed gas exchange, but kept the probe sterile via a Teflon membrane (MED-MEM, MED64). Samples were then transferred to the MED64 array system (MED64-Basic, MED64), and placed inside the head-plate of the array inside of an incubator (MCO-20AIC, Sanyo) to maintain CO2 levels for the culture. Cultures were allowed to equilibrate back to 5 % CO2 for 5 mins prior to initial recordings. Amplifiers were tuned and an initial playback was examined to determine whether probe was seated sufficiently in MEA system (Figure 2.7 C). Mobius (0.5.0, MED64) software was used to acquire recordings to .modat files. Input range was set to 2.3 mV (field standard for iPSC-derived neuron recordings) and data was acquired from all 64 channels were simultaneously. Traces were acquired for 15 min periods, or for 5 mins every 15 mins. Low cut frequency and high cut frequency was set to 0.1 Hz and 10000 Hz respectively to capture the full range available. To visualise raw data, it was exported as an ASCII text file. This file was then imported to Clampfit where it could be filtered to examine action potential, and generalised field potential changes.

## Results

To determine whether lower layer human cortical networks were able to exhibit oscillations, similar to those induced by muscarinic activation reported in anatomically intact rodent preparations ^28^, cultures were perfused with ACSF containing the non-specific cholinergic agonist carbachol (25 µM, CCh). In control conditions, networks exhibited either a lack of activity, sporadic firing with calcium elevations seen in single neurons or occasional synchronised bursts throughout the culture (Figure 1A). The addition of CCh resulted in a transformation of network activity. Within ∼60s networks exhibited waxing synchronised oscillations (Figure 1Ai-iv) with a regular periodicity (Figure 1Av, vi). This result was consistently and robustly seen in cultures tested (N >100). Synchronized oscillations could be sustained in the presence of CCh for over 30 minutes of imaging, maintaining their periodicity and frequency (Figure 1B). Oscillation periodicity exhibited a range across recordings with a median interval of 34.55 +/-17.85s (0.029 Hz, N = 100 cultures) (Figure 1B). There was no correlation of interval with culture days *in vitro* following attainment of functional maturity (Figure 1B). To determine role of singular neurons in manifestation of the observed oscillation, combined calcium imaging and patch clamp recordings were conducted. These revealed that oscillatory calcium increases were coincidental with neuronal depolarising events (Figure 1Ci; n = 5 preparations). These periodically instigated depolarisations in the presence of CCh differed from activity in control conditions and exhibited properties of UP and DOWN states ^26,33,41,42^, with depolarising states often crowned by trains of action potentials in frequency range 10 – 50 Hz (Figure 1Bii). Population LFP recording using multielectrode array and filtering for slow, low (< 1 Hz) and fast, high pass (> 100 Hz) filters revealed synchronised population activity across a 1 mm^2^ area (Figure 1D, inset), the events detected at a frequency of 0.036 Hz (N = 118 events) which were crowned by higher frequency neuronal firing at 54 Hz (N = 125 events) (Figure 1D).

**Figure 1.**
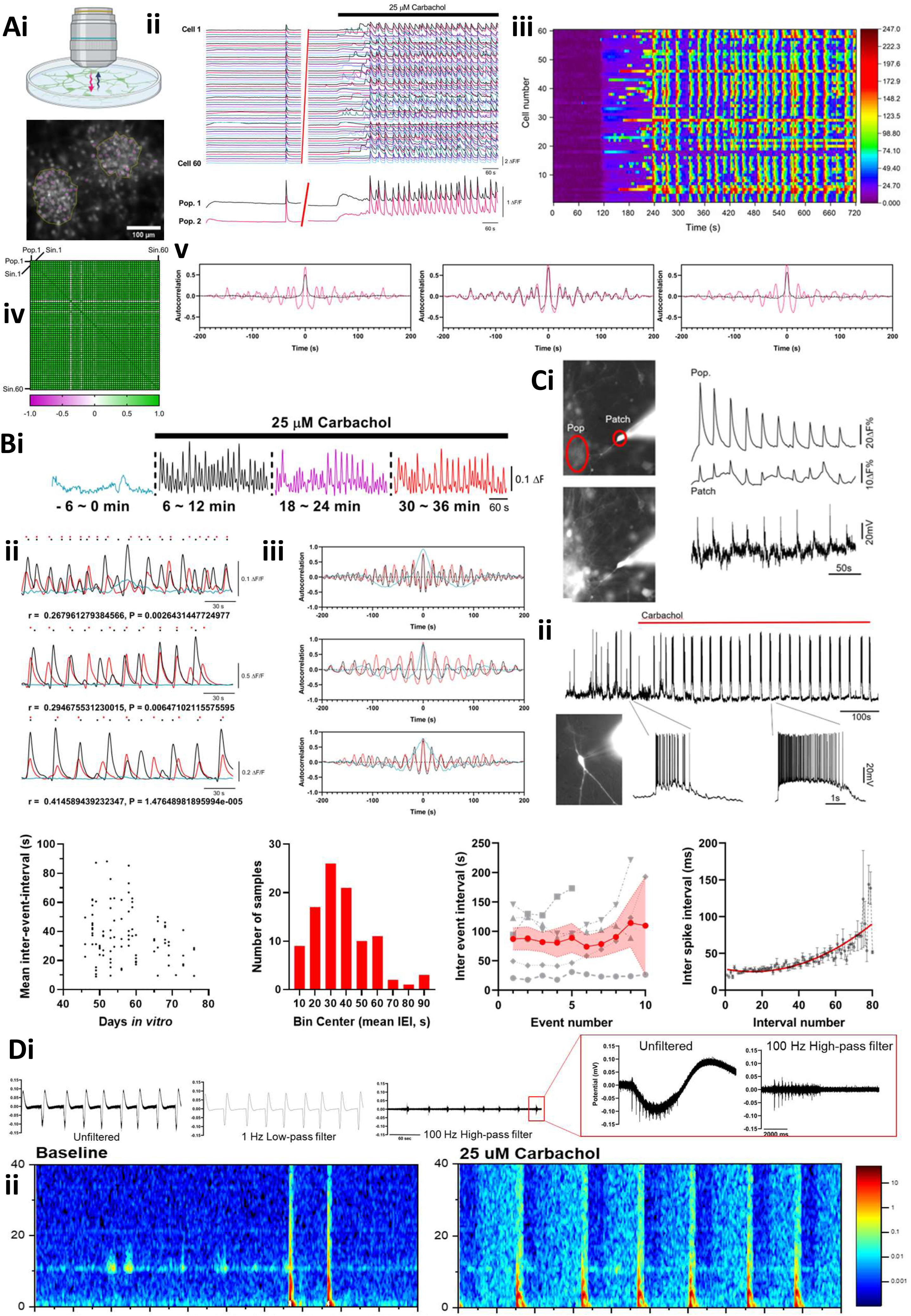
Carbachol induces periodic UP and DOWN state oscillations in human cortical networks. Ai) Live imaging of Fluo-4 AM calcium indicator allows detection of calcium dynamics. Representative field shows fluorescent intensity of Fluo-4 AM loaded hiPSC-derived co-cultures, yellow outlines denote regions of interest (ROIs) of two main cell populations and magenta outlines denote individual cells. (ii) Normalised fluorescence signal for single cell and population ROIs during baseline recording and post CCh (25uM, black bar) application. (iii) Raster plot displaying the relative fluorescence of active cells indicates periodic synchronisation in the presence of CCh. (iv) Correlation matrix plotting Pearson coefficient for cell populations 1-2 and individual cells 1-60 indicating synchronisation present via high correlation. (v). Autocorrelograms comparisons of right: single cell baseline (black) vs post CCh treatment (magenta), middle: single cell CCh treatment vs populational CCh treatment, and right: populational baseline (black) vs CCh treatment (magenta), illustrating periodicity of oscillations induced by CCh. (vi) Correlation matrix plotting Pearson coefficient for cell populations 1,2 and individual cells 1-60 indicating high correlation. Bi) Representative fluorescence trace of population activity from long-term imaging of extended CCh application. Traces display different time periods from a baseline prior to CCh addition, up to 36 minute CCh exposure. (-6-0mins cyan, 6-12mins black, 18-26 mins magenta, 30-36 mins red). (ii) Cross correlation of activity at baseline (cyan), 6-12 minutes (black) and 30-36 minutes (red) CCh exposure show sustained consistent periodicity of CCh induced oscillation. Digitalised signal intensity peaks represented in calcium release. N = 3 cultures. (iii) Autocorrelogram comparisons of baseline (cyan), 6-12 minutes (black) and 30-36 minutes (red) CCh, illustrating sustained oscillations induced by CCh. N = 3 cultures. Ci) Representative field images show simultaneous patch clamp recording from a neuron and calcium imaging during an oscillatory DOWN (top-left) and UP (bottom-left) states. Large red outline denotes ROI for neuronal population (Pop), whereas small red ROI encircles patch-clamped recorded neuron (Patch) for calcium analysis. Fluorescence traces show calcium changes in the cell population (top-right) and the patched neuron (middle-right). Voltage trace (bottom-right) from the simultaneous patch clamp recording, illustrating that calcium elevations are co-incidental with neuronal depolarisations. (ii) Representative example of a single neuron patch recording in current clamp mode, (inset) image of the neuron filled with Alexa488 dye. Upon CCh application, prior sporadic action potentials and burst firing (inset left) is transformed into periodic high-frequency burst firing (inset right) exhibiting UP and DOWN states. Di) Histogram displaying mean inter-event interval (IEI) of calcium release post-CCh addition over range of tested sample age. N = 100 cultures. (ii) Binned groupings of mean IEI post CCh addition. N = 100 cultures. (iii) Relationship of patch-clamp recorded inter-event intervals of UP states against event number for the first 10 events post carbachol addition for individual experiments (grey) with mean +/- sem (red). (iv) Relationship of patch-clamp recorded mean inter-spike intervals within UP states against interval number post carbachol addition for individual experiments with interpolated quadratic curve (red). Ei) Representative multi-electrode array recording of neuron and astrocyte co-cultures post CCh addition, displaying traces which are unfiltered (left), 1 Hz low-pass filtered for local field potential (middle), and 100 Hz high-pass filtered for individual spikes (right). Expanded (inset right), magnification of a single burst firing event showing local field potential deflection similar to UP state recordings both unfiltered and filtered for spikes. (ii) Representative spectrograms showing power spectrum density values of baseline (left) and post-CCh addition (right), highlighting periodicity and increased presence of high frequency activity.

**Figure 2.**
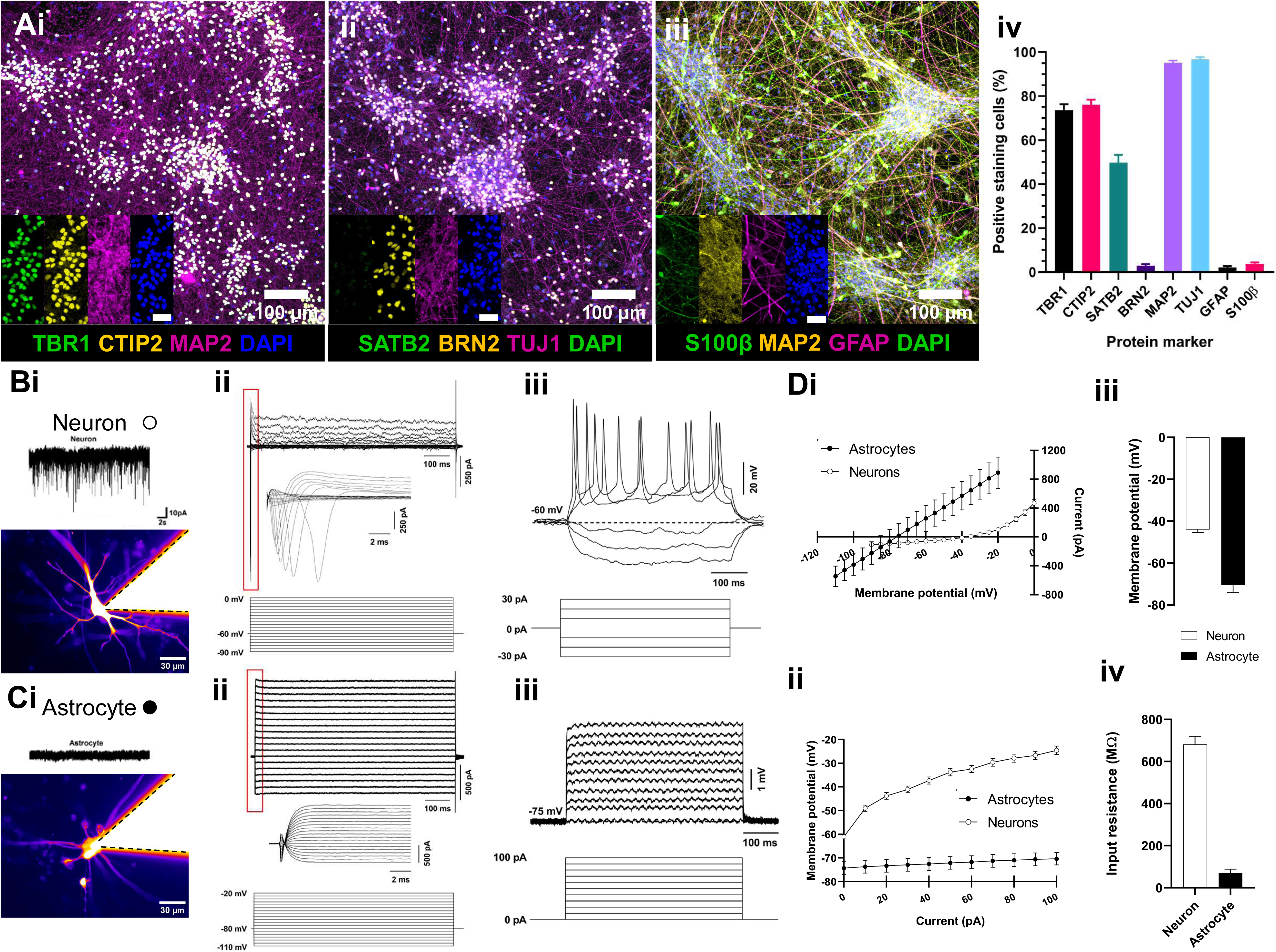
Cellular properties of differentiated lower layer cortical neurons and astrocytes. A) Representative images of immunostaining of, i) lower layer TBR1/CTIP2/MAP2+ neurons, ii) upper layer SATB2/BRN2/MAP2 neurons, iii) S100B/GFAP+ astrocytes and MAP2+ neurons. iv) Histogram showing the mean percentage of positively-expressing cells normalised by DAPI+ nuclei. N = 3 cultures. Bi) Representative image of Alexa488 filled neuron during whole cell patch clamp recording. ii) Representative recording from a neuron in voltage clamp, upper traces show current responses to voltage steps from -90mV to 0mV (step traces below) from a holding potential of - 60mV. Inset traces expand first 10ms following depolarising step illustrating fast inward sodium currents. iii) Representative recording from a neuron in current clamp mode, and responses to current steps from -30 to +30pA. More positive steps elicit action potential firing. Ci) Representative image of Alexa488 filled astrocyte during whole cell patch clamp recording. ii) Representative recording from a astrocyte in voltage clamp, upper traces show current responses to voltage steps from -110mV to -20mV (step traces below) from a holding potential of -80mV. Inset traces expand first 10ms following depolarising steps. iii) Representative recording from an astrocyte in current clamp mode, and responses to current steps from 0 to +100pA. Di) Current-voltage plot from the example neuron in B (white) and astrocyte in C (black). ii) Voltage-Current plot for the same cells. Plots showing iii) input resistance (N= 101 neurons, 13 astrocytes) and iv) resting membrane potential values (N=47 neurons, 4 astrocytes) for neurons and astrocytes.

To confirm cellular identity and functionality of generated networks, cultures were immunostained for protein expression, and single cells interrogated electrophysiologically. Immunocytochemical staining of electrophysiologically-mature cultures showed highly positive expression of expected neuronal markers CTIP2, TBR1, SATB2, TUJ1 and MAP2 (Figure 2A, 2B) and lower proportions of BRN2, and astrocytic GFAP and S100b expression, consistent with differentiated layer 5/6 cortical neurons and supporting levels of astrocytes, with very few layer 2/3 neurons. Filling of single neurons with Alexa 488 fluorescent dyes revealed typical neuronal morphological properties with 14.22± 0.47 µm (n = 51) soma diameter and 2-4 primary dendrites (Figure 2B). Neurons exhibited voltage-dependent ionic currents such as fast inward Na^+^ currents on depolarisation, and action potential firing to increased depolarisation in current clamp mode (Figure 2B). Astrocytes filled with fluorescent dyes revealed large cell soma diameters (23.45 ± 1.72 µm, n=15, Figure 2C) and long processes, with some evident gap-junctional coupling from filling of neighbouring astrocytes (Figure 2C). Electrophysiological properties were consistent with respective cell identity, with astrocytes exhibiting lower input resistance (neurons: 681 ± 39.9MΩ, n=47; astrocytes: 69.6 ± 18.62MΩ, n=4) more negative membrane potential (neurons: -43.9 ± 1.39 mV, n=47; astrocytes: -70.5 ± 3.38 mV, n=4), linear current voltage relationships lack of action potential firing.

To determine whether the observed oscillatory properties of the networks were a result of specific cholinergic-induced effect or an inherent network property induced by an increase in activity, we increased extracellular potassium, from 2.5 to 7.5 mM [K^+^Cl^−^]_O_ (K-aCSF), to depolarise the neurons and increase network excitability. Increased general excitability was seen with an increase in synchronised events across the cell populations (Figure 3A). Activity however lacked the regularity of CCh-induced oscillations (Figure 3Av), and was not sustained in the same way. Single neuron patch clamp recordings revealed that K-aCSF perfusion lead to an increase in general synaptic activity and neuronal depolarisation often leading to synchronisation (Fig.3B). Depolarising events in K-aCSF however did not exhibit features of UP and DOWN states, they had lower amplitude (CCh: 18.74 ± 1.6mV, n=13, K-aCSF:10.00 ± 1.67, n=6, P<0.01), shorter inter-burst-event interval (CCh: 26.09 ± 2.83s, n=13, K-aCSF: 13.38 ± 0.86, n=6, P<0.01) translating as higher oscillation frequency (CCh:0.04 ± 0.0043Hz, n=13, K-aCSF: 0.076 ± 0.0045, n=6, P<0.01) and importantly time between events of greater variance than the UP states induced in CCh (CCh CV^2^: 17.16 ± 3.2, K-aCSF CV^2^: 47.03 ± 6.87) (Figure 3C). Expression of synapsin-ChR2 in neurons and its activation at a similar slow oscillation rate did not result in induction of an ongoing oscillation indicating that the oscillation requires an ongoing drive such as the action of CCh (Figure 3E). Together the results show that CCh induced slow oscillatory periodic activity that represents a specific mode of activity.

**Figure 3.**
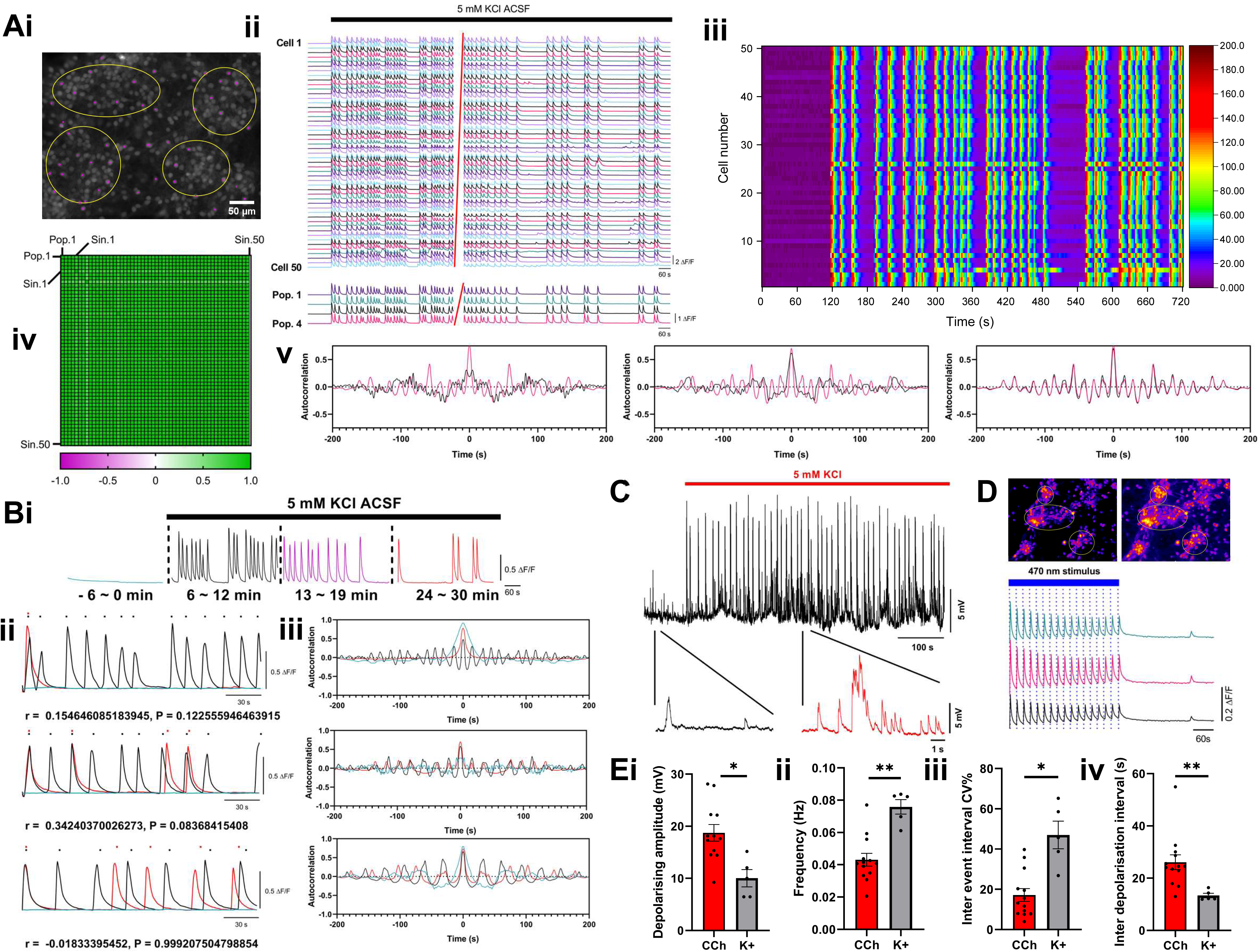
Network activity induced by elevated potassium is distinct from that induced by CCh. Ai) Representative field shows fluorescent intensity of Fluo-4 AM loaded hiPSC-derived co-cultures, yellow outlines denote regions of interest (ROIs) of two main cell populations and magenta outlines denote individual cells. (ii) Normalised fluorescence signal for single cell and population ROIs during immediate and prolonged elevated KCl ACSF (5mM, black bar) application.(iii) Raster plot displaying the relative fluorescence of active cells indicates periodic synchronisation in the presence of KCl-ACSF. (iv) Correlation matrix plotting Pearson coefficient for cell populations 1-4 and individual cells 1-50 indicating synchronisation present via high correlation. (v). Autocorrelograms comparisons of right: single cell baseline (black) vs post CCh treatment (magenta), middle: populational baseline (black) vs CCh treatment (magenta), and right: single cell CCh treatment (magenta) vs populational CCh treatment (black) illustrating periodicity of oscillations induced by CCh. Bi) Representative fluorescence trace of population activity from long-term imaging of extended elevated KCl ACSF application. Traces display different time periods from a baseline prior to CCh addition, up to 30 minute KCl-ACSF exposure. (-6-0mins cyan, 6-12mins black, 13-19 mins magenta, 24-30 mins red). (ii) Cross correlation of activity at baseline (cyan), 6-12 minutes (black) and 24-30 minutes (red) KCl-ACSF exposure showing a lack of sustained consistent periodicity of elevated KCl-induced oscillation. Digitalised signal intensity peaks represented in calcium release. N = 3 cultures. Autocorrelogram comparisons of baseline (cyan), 6-12 minutes (black) and 24-30 minutes (red) KCl-ACSF addition, illustrating inconsistent burst firing profiles induced by KCl. N = 3 cultures. C) Representative example of a single neuron patch recording in current clamp mode after application of elevated KCl-ACSF, (inset) expanded view of pre and post application. Upon KCl-ACSF application, prior sporadic action potentials (inset left, black) is transformed into higher frequency action potentials and large depolarising events (inset right, red). D) Representative field images pre and post stimulation of Chr2 optogenetic constructs expressed in neurons. Representative traces show elevation in calcium in Rhod-2AM filled cultures that coincide with 470nm stimulus (1000ms) every 20 seconds. E) Histograms comparing patch-clamp single neuron properties post CCh (red) or KCl-ACSF (grey) addition, including (i) depolarising amplitude, (ii) inter-event-interval frequency, (iii) inter-event-interval coefficient of variance, and iv) length of inter depolarisation interval. Mean +/-SEM.

To determine the signalling mediators of network activity during the slow oscillation, we used immunocytochemical characterisation and interventions. Immunocytochemical (ICC) staining was conducted for protein markers hypothesised to be present based upon UP-state and slow oscillations in rodent preparations ^28^. Muscarinic M1 and M3 receptors were expressed on neurons and astroctyres, whereas vesicular glutamate transporter VGlut1 was expressed in neurons and not astrocytes. Staining for the GABAergic neuron marker GAD67 was observed, however there was no identifiable somatic staining, which is the usual criteria for acceptance of GABAergic neuron presence. To determine the neurotransmitter receptors involved in the instigation and sustainment of the CCh instigated slow oscillation we used population calcium imaging in conjunction with pharmacological interventions. CCh-induced oscillations in mouse cortex have previously been shown to act via cholinergic muscarinic receptors, but not nicotinic receptors ^43^. We found that selective muscarinic agonist oxotremorine induced oscillations similar to the observed CCh-induced oscillations (0.86 ± 0.075/min, n=9/9 experiments). Whilst M1 and M3 muscarinic receptor subtype antagonists telenzipine (CCh: 1.94 ± 0.361, CCh+TLZ: 0.21 ± 0.03, P=0.00134, n=5) and 4DAMP (CCh: 1.45 ± 0.26, CCh+4-DAMP: 0.26 ± 0.07, P=0.0051, n=4) silenced ongoing CCh-induced oscillations (Fig.4.B). The results therefore indicate that M1/M3 receptors, and their downstream GPCR subunit activation is necessary and sufficient to induce CCh-induced oscillatory UP state activity in human cortical excitatory networks.

**Figure 4.**
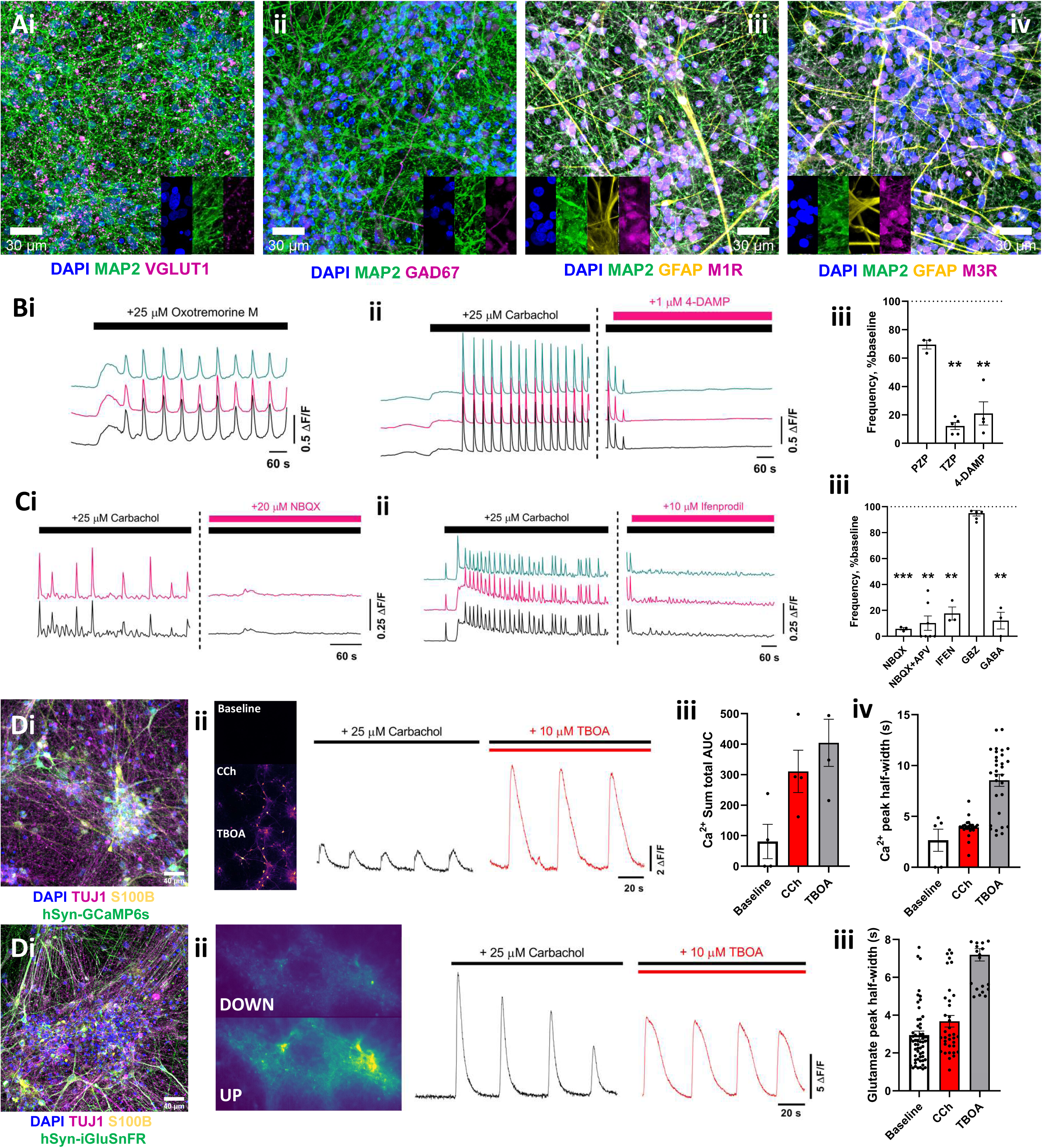
The muscarinic network oscillactivity is glutamate driven and controlled by EAATs. Immunocytochemical staining showing cellular expression patterns of vesicular glutamate transporter VGlut1, glutamate decarboxylase GAD67, and muscarinic M_1_ and M_3_ G_q_-coupled receptors (colour key under each image). B) Representative fluorescence traces showing instigation of oscillation by (i) Oxotremorine M (25 µM, black bar), and (ii) block of CCh-induced oscillation by 4-DAMP (1 µM, magenta bar). (iii) Bargraph shows normalised summary data of effect of m1 (Pirenzipine and Telenzipine) and m3 (4-DAMP) selective antagonists on oscillation up-state frequency. C) Representative fluorescence traces illustrating the effect of (i) AMPAR antagonist NBQX and (ii) NMDAR antagonist Ifenprodil. (iii) Bargraph shows normalised summary data of effect of neurotransmitter receptor antagonists and agonist GABA on CCh oscillation UP-state frequency. Di) Immunocytochemical staining showing cellular expression patterns of calcium indicator hSyn-GCAMP6s genetic construct. (ii) Representative fields showing relative fluorescence increase in GCAMP6s construct under different treatment conditions, and fluorescence traces from the same experiment. Each trace is an average of 10 neurons during CCh-induced oscillation and following further application of TBOA. Summary bargraphs show (iii) total area under the curve, and pooled half widths of UP-state events for different conditions across a N=4 experiments. Ei) Immunocytochemical staining showing cellular expression patterns of glutamate indicator hSyn-iGluSNFR genetic construct. (ii) Representative fields showing relative fluorescence increase in iGluSNFR construct during a DOWN and UP-state, and fluorescent traces display the iGluSNFR fluorescence during CCh-induced oscillation and following further application of TBOA. (iii) Bargraph shows pooled half widths of UP-state events for different conditions across a N=4 experiments.

Because the differentiation protocols employed were aimed at producing excitatory lower layer cortical neurons and the main receptors mediating cortical synaptic transmission are AMPA and NMDA receptors we tested the effect of selective antagonists to these respectively. Application of AMPA receptor antagonist NBQX (20 µM) caused a reduction in population network activity (CCh:2.16 ± 0.386, CCH+NBQX: 0.115 ± 0.004, P=0.0056 n=3), as did NMDA receptor antagonist APV (CCh: 2.36 ± 0.37, CCh+NBQX+APV:0.143 ± 0.076, P=8×10^−5^, n=5), and NR2B containing receptor antagonist Ifenprodil (CCh: 2.47 ± 0.45, CCh+Ifen: 0.461 ± 0.191, P=0.015, n= 3; Figure 4C). These results indicate that sustainment of the CCh-induced oscillation is dependent on ionotropic glutamate receptor mediated synaptic transmission. The main inhibitory transmitter in the brain is GABA, which is involved in the generation of several types of oscillatory activity ^44^. However, application of the GABA-A antagonist Gabazine (20 µM) had no significant effect on CCh-induced oscillations (CCh 1.77 ± 0.102, CCh+GBZ: 1.68 ± 0.098, P=0.54, n=6) (Fig.4C). This is expected since no inhibitory GABAergic neurons would be expected in the excitatory cortical directed cultures. Application of GABA however caused a significant reduction in activity (CCh: 1.679 ± 0.16, CCh+GABA: 0.206 ± 0.104, P=0.0015, n=3) indicating that whilst there may be no intrinsic GABAergic signalling in the networks, consistent with the ICC findings (Figure 4A), the excitatory neurons *are* expressing functional GABA receptors. To determine whether the glutamate synaptic transmission during the slow oscillation was controlled by EAAT glutamate transporters we added the non-specific GLT1/GLAST antagonist TBOA. The inhibitor effected a change in the CCh-induced oscillation indicating that EAAT glutamate transporters are important functional elements controlling synaptic transmission in the monolayer human cortical cultures (Figure 4D). Changes in the kinetic profile of the calcium signals during TBOA were clearly evident. Immediate responses exhibited increased amplitude, while duration of UP-state related calcium signals also increased. (CCh event half-width: 3.9 ± 0.16s, TBOA event half-width: 8.55 ± 0.59, P=1.74×10-9, n=27,30 events, 4 experiments). However, the degree of photobleaching of GCaMP6 during extended high-speed acquisition made determination of changes in amplitude and areas difficult (Figure 4D). To attempt to determine whether EEAT inhibition resulted in the increased glutamate in the networks, we utilised the genetically encoded glutamate indicator iGluSNFr. Using the human synapsin promoter, expression was targeted to neurons (Figure 4E). iGluSNFr imaging in the presence of CCh revealed a slow oscillation pattern in glutamate release, indicating neuronal network synaptic synchronisation. In the presence of TBOA, the oscillatory pattern was retained, with the glutamate signal increased significantly (Figure 4F; CCh event half-width:3.67 ± 0.31s, TBOA event half-width: 7.19 ± 0.33s, P=1.87×10-11, n=46, 25 events, 3 experiments). Confirming that glutamate transmission and glutamate signalling is controlled by EAATs during the CCh induced oscillation. Taken together therefore, the results show that CCh-induced UP and DOWN states are sustained by synaptic glutamate release ^34,43^, and that fidelity of the signalling is controlled by glutamate transporters.

In rodent and cat preparations purinergic signalling has been shown to have roles in setting the periodicity of neuronal oscillations ^35^. A possible mediator released in response to neuronal activity is Adenosine, which by its action at presynaptic A1 receptors could periodically inhibit excitation ^34,43^. The cultures showed that A1R receptors were present in both neurons and astrocytes (Figure 5A). In these hiPSC-derived layer 5/6 networks, block of A1 receptors by the antagonist 8-CPT indeed disrupted the regular periodicity of the slow waves and reduced CCh induced network synchronisation. This was seen in calcium imaging recordings by acquiring neuronal population level fluorescence (Figure 5B; CCh:12.07 ± 0.77, 8-CPT: 8.18 ± 0.75, n=3, P=0.0199).

**Figure 5.**
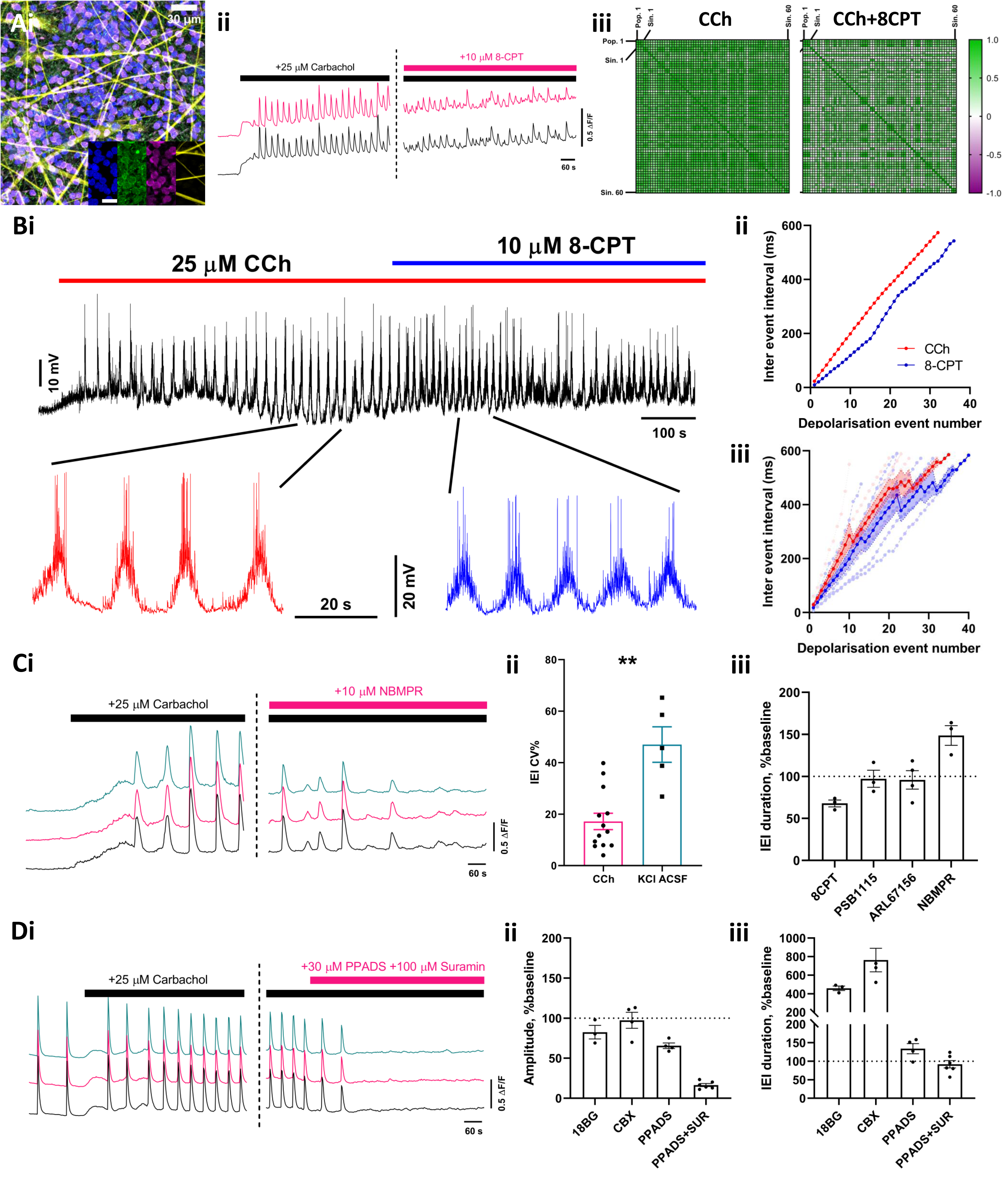
CCh induced oscillation periodicity is controlled by Adenosine A1 receptors. Ai) Immunocytochemical staining showing cellular expression patterns of adenosine A_1_ receptor staining (magenta) with neuronal (MAP2, green) and astrocyte (GFAP, yellow) protein markers. (ii) Fluorescence traces from population ROIs showing the effect of A_1_R antagonist 8-CPT on CCh-induced oscillation. (iii) Cross-correlation representations display degree of correlation between single cells during CCh-induced oscillation and in the presence of additional 8-CPT. Bi) Example single neuron patch clamp recording in current clamp where following oscillation induction by CCh application and following A_1_R antagonist 8-CPT application. Expanded traces (inset) show 60s sections of activity in application of CCh (red) and CCh with 8-CPT (blue). (ii) Plot shows normalised IEI time against event number for the cell shown in the example trace, exhibiting a linear relationship in CCh which is disrupted and shallower in the presence of 8-CPT. (iii) Results from all recorded cells are shown, with the mean per depolarisation event shown in bold, and the standard deviation shown by the coloured outline. Ci) Representative traces of population ROIs illustrating the effect of ENT1 inhibitor NMBPR on CCh induced slow oscillation. (ii) Bargraph representing the coefficience of variance in CCh and KCl-induced oscillations. (iii) Summary bargraph displaying effect of different pharmacological interventions to nucleoside equilibrium in slow oscillation periodicity, normalised to control CCh IEI. Di) Representative traces of population ROIs illustrating the effect of purinergic inhibitors PPADS and Suramin on CCh induced slow oscillation. (ii) Summary bargraph displaying effect of different pharmacological interventions to gap junctions and purinergic signalling in slow oscillation periodicity, normalised to control CCh IEI.

In single neuron electrophysiological recordings this disruption was also evident, resulting in an overall decrease in inter-burst-interval in 4 recordings and a small increase in 2 (CCh: 22.73 ± 1.81s, 8-CPT: 20.39 ± 2.2s, n=6, P=0.14). A decrease in action potential firing of ∼50% following 8-CPT was seen, indicating a reduction in excitability likely a result of synchronisation disruption leading to reduced excitation coherence (CCh: 160 ± 28 APs in 5mins, 8CPT: 95 ± 26.76, P<0.05, n=6) . In contrast the A3a antagonist PSB1115 had no effect on IEI (CCh:68 ± 9.17, PSB: 66.02 ± 8.37, n=3, P=0.76). ATP is metabolised to ADP and adenosine which can act as a source of signalling adenosine. To gain insight into the source of adenosine, ectonucleotidase inhibitor ARL 67156 was applied. There was no change in oscillation frequency (CCh: 17.29 ± 4.01, ARL: 15.67 ± 3.2, n=3, P=0.517) nor amplitude, indicating that ATP was not the source of the Adenosine in the cultures. Another important component of adenosine homeostasis is equilibrative adenosine transporters (ENTs). To this end, the ENTs inhibitor NMPBR resulted in similar oscillation disruption to that caused by 8-CPT (CCh: 37.089 ± 0.17, NBMPR: 55.21, n=3, P=0.054) suggesting that ENTs control of adenosine levels are important in maintaining the oscillation and overall activity control. Adenosine can be released or taken up via these transporters. So results are consistent with adenosine acting via A1 receptors controlling oscillation periodicity.

Immunocytochemical staining for muscarinic receptors expression revealed expression on both neurons and astrocytes, and similarly for the purinergic A1 receptor (Figure 4a). In bath application experiments astrocytes seem to respond to CCh with a large calcium elevations before neurons, and prior to network oscillation instigation. In the absence of neurons in astrocyte monoculture, astrocytes exhibited occasional and sporadic spontaneous calcium elevations and exhibited robust, non-synchronised calcium oscillations in response to CCh addition (Figure 6A). In contrast, there was no difference in astrocyte calcium activity in control aCSF and K-aCSF, indicating a specific astrocyte muscarininc receptor response to CCh. However, to identify the specific individual cell type contribution of activity during CCh in co-cultures, neuron-and astrocyte-expressing GECI cultures were prepared separately. In neuronal cultures cells remained highly synchronised, and all activity decreased following incubation with TTX, indicating a dependence on action potential driven activity. However, astrocytes displayed non-synchronised activity, which whilst decreased with TTX, was not fully extinguished, indicating a contribution of neuronal neurotransmitter release to astrocyte calcium activity. To determine a potential role for astrocyte calcium signalling in slow wave oscillations, we targeted astrocyte calcium signalling using BAPTA intracellular infusion to buffer intracellular astrocyte calcium. Cultures were prepared using GCaMP6 expressing neurons, and astrocytes not expressing any constructs. Visually identified astrocytes in monolayer cultures were patch clamped with pipettes containing BAPTA internal solution. Following infusion, CCh-induced slow oscillation activity was recorded and compared to activity in a different region of the same 2D culture where astrocytes were not BAPTA filled. Inclusion of Alexa 550 in the patch pipette enabled monitoring of cell filling and occasional filling of other astrocytes, presumably via gap junctions (Figure 6C). Slow oscillation activity acquired from areas where astrocyte calcium was buffered displayed shorter intervals between UP state calcium elevations (CCh: 71.28 ± 6.78s, BAPTA: 40.11 ± 2.28s, P< 0.01, n=4) and hence an increased oscillation frequency (CCh: 0.014 ± 0.0016Hz, BAPTA: 0.025 ± 0.0014, P< 0.05, n=4). To support this hypothesis, gap junction blocking was also undertaken in cocultures, which similarly altered or silenced ongoing CCh-induced oscilations, strengthening the likelihood of astrocytic involvement in oscillation.

**Figure 6.**
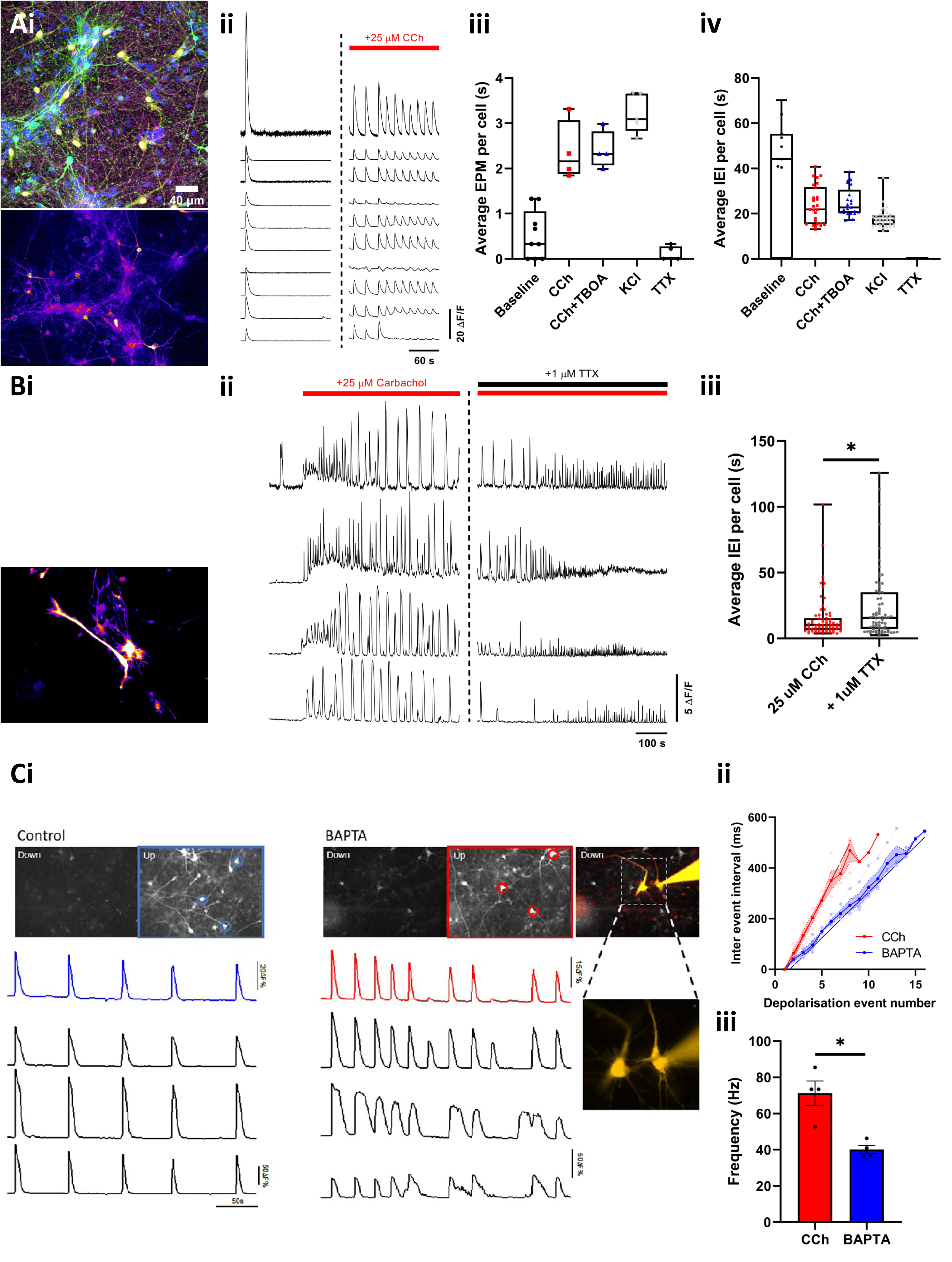
Neuronal oscillation periodicity is dependent on astrocyte calcium signalling. Ai) Immunocytochemical staining showing cellular expression patterns of calcium indicator hSyn-GCAMP6s genetic construct and representative field showing relative fluorescence increase in GCAMP6s construct after treatment with CCh. (ii) Representative fluorescence traces from high-speed imaging of the GCAMP6s construct in individual neurons, showing similar trace profiles to Fluo4 imaging of CCh-induced oscillations. Summary bargraphs displaying effect of different pharmacological interventions to (iii) UP-state frequency and (iv) event interval. Bi) Immunocytochemical staining showing cellular expression patterns of calcium indicator GFAP-GCAMP6f genetic construct and representative field showing relative fluorescence increase in GCAMP6f construct after treatment with CCh. (ii) Representative fluorescence traces from high-speed imaging of the GCAMP6f construct in individual astrocytes, showing continued, albeit altered, oscillatory activity after TTX addition compared to loss of activity in neuronal CCh-induced oscillations. (iii) Bargraph quantifying inter-event-interval after CCh, and subsequent TTX addition, showing activity independently of neuronal action potentials. Ci) Effect of BAPTA chelation of calcium in patch-clamped astrocytic syncytium. Top: representative field of views of neurons expressing hSyn-GCaMP6s in the presence of CCh or CCh and BAPTA addition, leftmost image shows cells during a DOWN-state and image on right field during an UP-state. Fluorescence changes of whole field population (denoted by blue or red border ROI), and individually ringed neurons are plotted below. Representative images on the right also have combined image of astrocyte patch clamped with an internal solution containing calcium chelator BAPTA and Alexa 555 for visualisation. Inset: patching of a single astrocyte also results in the filling of a neighbouring astrocyte. (ii) Plot showing normalised IEI time against event number for pairs of recordings from the same coverslips in regions without BAPTA infusion (red) and in those where astrocytes were infused with BAPTA (blue) showing a shallower relationship when astrocyte calcium signalling is buffered by BAPTA infusion. (iii) Bargraph showing changes in firing frequency of patched neurons recorded in the same experiments, showing reduced neuronal firing in BAPTA-infused astrocyte conditions.

After establishing a co-dependency of neurons and astrocytes for oscillation maintenance, we next aimed to determine whether neurons or astrocytes were the key cellular initiators. For this we used a chemogenetic approach. From insights gained from pharmacological interventions we used AAV transduction with a construct containing the Gq coupled human M3 receptor construct hM3Dq, targeting expression to either neurons or astrocytes using specific promoters. To confirm the functionality of this approach to activate the Gq-IP3R pathway we first used astrocyte monocultures. Application of the DREADD agonist CNO resulted in Ca^2+^ elevations in individual astrocytes that exhibited a range of activity patterns similar to CCh stimulation, demonstrating the efficacy of this approach (Figure 7A).

**Figure 7.**
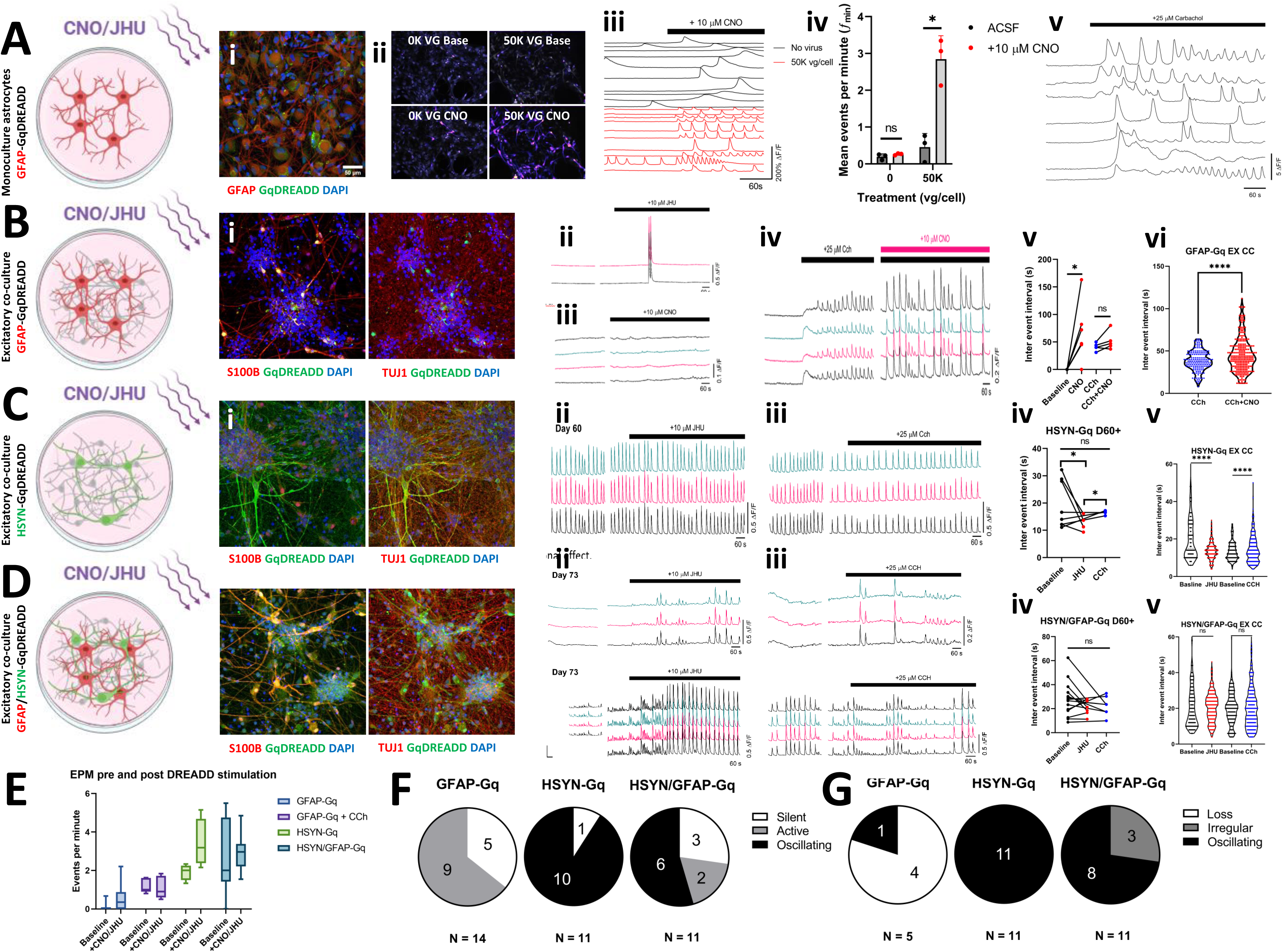
Neuronal muscarinic Gq-coupled receptor activation is necessary and sufficient to induce slow oscillations. Monoculture of human iPSC-derived astrocytes transfected with GFAP-hM3Dq DREADD. (i) Immunocytochemical staining showing cellular expression patterns of DREADD hM3Dq construct in astrocytes (GFAP). (ii) Representative field showing relative fluorescence increase in Fluo4 AM-loaded astrocytes treated with 0 viral genomes and 50,000 viral genomes of GFAP-hM3Dq after addition of DREADD agonist CNO. (iii) Representative fluorescence traces from calcium imaging in individual astrocytes expressing DREADDs or control astrocytes during addition of DREADD agonist CNO. (iv) Average events per minute in astrocytes expressing DREADDs or control astrocytes before and after addition of DREADD agonist CNO. (v) Representative fluorescence traces from calcium imaging in individual astrocytes after addition of CCh. B) Co-culture of human iPSC-derived excitatory neurons and astrocytes transfected with GFAP-hM3Dq DREADD. (i) Immunocytochemical staining showing cellular expression patterns of DREADD hM3Dq construct in astrocytes (GFAP) and neurons (TUJ1).

To test the hypothesis that astrocytes were the initiators of slow oscillations, we combined hM3Dq expressing astrocytes with non-transduced cortical neurons in co-culture. Activation of the DREADD receptor with CNO or JHU in the absence of CCh did not result in a neuronal slow oscillation (Fig 7B), indicating that astrocyte muscarinic activation is insufficient for network oscillation generation. When slow oscillations were induced in the cocultures with CCh however, and astrocyte-specific Gq DREADD receptors stimulated subsequently, the oscillation became less regulated, similar to the effect of 8CPT. When oscillations were induced by CCh, however, and astrocytes then activated by DREADD agonist, the result was a dysregulation of the CCh oscillation periodicity and a lengthening of inter event interval (Fig.7).

In experiments where only neurons expressed the hM3Dq construct, stimulation with DREADD agonists resulted in instigation of a slow oscillation (Figure 7C), unlike the stimulation of cultures where astrocytes expressed the construct. Indeed, in co-cultures where the hM3Dq DREADD was expressed in neurons, we found that cultures expressed spontaneous slow oscillations. Agonist activation of the neuronal DREADD resulted in non-significant, with addition of CCh to activate the native muscarinic pathway in neurons and astrocytes resulted in a slowing of the oscillation (Fig 7).

Expressing the DREADD in both astrocytes and neurons produced a similar situation to that for selective neuronal expression, in that cultures displayed spontaneous slow oscillations that in instances were strengthened by DREADD agonist and slowed by CCh activation of muscarinic receptors. However, the result was inconsistent and ultimately non-significant, indicating a conflicting action of both neurons and astrocytes. Taken together, the results show that activation of neuronal Gq-IP3 receptor mediated pathways is sufficient and necessary to induce a slow network oscillation.

## Discussion

Human iPSCs have allowed the differentiation of cell models that resemble a wide variety of CNS neurons on a molecular and genetic level – however it remains uncertain to what extent iPSC-derived neural cells can generate high-order functional activity similar to that of *ex vivo* tissue. To establish this we require network models that resemble known phenomena of neural activity.

In this study we found that selective muscarinic receptor activation instigated a slow (< 1 Hz) oscillation with regular, long-term periodicity in networks of hiPSC-derived cortical layer V/VI neuron and astrocyte cultures. The activity was distinct from that induced by ↑[K+]_O_ lowering the depolarisation threshold, and thus reflects the unique effect of ascending neuromodulatory cholinergic signalling in lower layer cortical neurons. The slow oscillation was defined by periodic, glutamatergic synapse-induced neuronal depolarisations with the general features of UP states. Oscillation maintenance showed mechanistic co-dependence on purinergic signalling, and its periodicity was disrupted by A_1_ receptor agonists, blockage of gap junctions, and calcium chelation in local astrocytes. However, it was not dependent on a GABAergic tone to facilitate inhibition. Thus, periodicity is dependent on astrocyte calcium signalling, although neuronal activation is sufficient and necessary for the instigation of the oscillation.

## Conclusion

These findings indicate that differentiated hiPSC-derived cortical lower layer neurons have the inherent capacity to generate this activity and to respond to the physiological cholinergic tonic drive. Whether this mechanism represents a key driver exclusively during development, or serves a key function in adult brain activity remains unknown. However, it is clear with further investigation and development of improved models which incorporate a more complex network structure, we may be able to faithfully recapitulate mechanisms of physiological activity in vitro, which would aide in the identification of abnormalities to oscillatory mechanisms in disease states.

## Notes

### Competing Interest Statement

The authors have declared no competing interest.

## Bibliography

1. Cebolla, A.M., and Cheron, G. (2019). Understanding Neural Oscillations in the Human Brain: From Movement to Consciousness and Vice Versa. Front. Psychol. 10. 10.3389/fpsyg.2019.01930.

2. Buzsáki, G., and Draguhn, A. (2004). Neuronal Oscillations in Cortical Networks. Science 304, 1926–1929. 10.1126/science.1099745.

3. Little, S., and Brown, P. (2014). The functional role of beta oscillations in Parkinson’s disease. Parkinsonism Relat. Disord. 20, S44–S48. 10.1016/S1353-8020(13)70013-0.

4. Cole, S.R., Meij, R. van der, Peterson, E.J., Hemptinne, C. de, Starr, P.A., and Voytek, B. (2017). Nonsinusoidal Beta Oscillations Reflect Cortical Pathophysiology in Parkinson’s Disease. J. Neurosci. 37, 4830. 10.1523/JNEUROSCI.2208-16.2017.

5. McCarthy, M.M., Moore-Kochlacs, C., Gu, X., Boyden, E.S., Han, X., and Kopell, N. (2011). Striatal origin of the pathologic beta oscillations in Parkinson’s disease. Proc. Natl. Acad. Sci. 108, 11620–11625. 10.1073/pnas.1107748108.

6. Osipova, D., Ahveninen, J., Jensen, O., Ylikoski, A., and Pekkonen, E. (2005). Altered generation of spontaneous oscillations in Alzheimer’s disease. NeuroImage 27, 835–841. 10.1016/j.neuroimage.2005.05.011.

7. Murty, D.V., Manikandan, K., Kumar, W.S., Ramesh, R.G., Purokayastha, S., Nagendra, B., Ml, A., Balakrishnan, A., Javali, M., Rao, N.P., et al. (2021). Stimulus-induced gamma rhythms are weaker in human elderly with mild cognitive impairment and Alzheimer’s disease. eLife 10, e61666. 10.7554/eLife.61666.

8. Benedek, K., Berényi, A., Gombkötő, P., Piilgaard, H., and Lauritzen, M. (2016). Neocortical gamma oscillations in idiopathic generalized epilepsy. Epilepsia 57, 796–804. 10.1111/epi.13355.

9. Louis, E.K.S., Frey, L.C., Britton, J.W., Frey, L.C., Hopp, J.L., Korb, P., Koubeissi, M.Z., Lievens, W.E., Pestana-Knight, E.M., and Louis, E.K.S. (2016). The Developmental EEG: Premature, Neonatal, Infant, and Children. In Electroencephalography (EEG): An Introductory Text and Atlas of Normal and Abnormal Findings in Adults, Children, and Infants [Internet] (American Epilepsy Society).

10. Ibarra-Lecue, I., Haegens, S., and Harris, A.Z. (2022). Breaking Down a Rhythm: Dissecting the Mechanisms Underlying Task-Related Neural Oscillations. Front. Neural Circuits 16. 10.3389/fncir.2022.846905.

11. Buzsáki, G., and Wang, X.-J. (2012). Mechanisms of Gamma Oscillations. Annu. Rev. Neurosci. 35, 203. 10.1146/annurev-neuro-062111-150444.

12. Oberheim, N.A., Takano, T., Han, X., He, W., Lin, J.H.C., Wang, F., Xu, Q., Wyatt, J.D., Pilcher, W., Ojemann, J.G., et al. (2009). Uniquely Hominid Features of Adult Human Astrocytes. J. Neurosci. 29, 3276–3287. 10.1523/JNEUROSCI.4707-08.2009.

13. Li, J., Pan, L., Pembroke, W.G., Rexach, J.E., Godoy, M.I., Condro, M.C., Alvarado, A.G., Harteni, M., Chen, Y.-W., Stiles, L., et al. (2021). Conservation and divergence of vulnerability and responses to stressors between human and mouse astrocytes. Nat. Commun. 12, 3958. 10.1038/s41467-021-24232-3.

14. Chambers, S.M., Fasano, C.A., Papapetrou, E.P., Tomishima, M., Sadelain, M., and Studer, L. (2009). Highly efficient neural conversion of human ES and iPS cells by dual inhibition of SMAD signaling. Nat. Biotechnol. 27, 275–280. 10.1038/nbt.1529.

15. Shi, Y., Kirwan, P., and Livesey, F.J. (2012). Directed differentiation of human pluripotent stem cells to cerebral cortex neurons and neural networks. Nat. Protoc. 7, 1836–1846. 10.1038/nprot.2012.116.

16. Bardy, C., van den Hurk, M., Eames, T., Marchand, C., Hernandez, R.V., Kellogg, M., Gorris, M., Galet, B., Palomares, V., Brown, J., et al. (2015). Neuronal medium that supports basic synaptic functions and activity of human neurons in vitro. Proc. Natl. Acad. Sci. 112, E2725–E2734. 10.1073/pnas.1504393112.

17. Bardy, C., van den Hurk, M., Kakaradov, B., Erwin, J.A., Jaeger, B.N., Hernandez, R.V., Eames, T., Paucar, A.A., Gorris, M., Marchand, C., et al. (2016). Predicting the functional states of human iPSC-derived neurons with single-cell RNA-seq and electrophysiology. Mol. Psychiatry 21, 1573–1588. 10.1038/mp.2016.158.

18. Trujillo, C.A., Gao, R., Negraes, P.D., Gu, J., Buchanan, J., Preissl, S., Wang, A., Wu, W., Haddad, G.G., Chaim, I.A., et al. (2019). Complex Oscillatory Waves Emerging from Cortical Organoids Model Early Human Brain Network Development. Cell Stem Cell 25, 558–569.e7. 10.1016/j.stem.2019.08.002.

19. Samarasinghe, R.A., Miranda, O.A., Buth, J.E., Mitchell, S., Ferando, I., Watanabe, M., Allison, T.F., Kurdian, A., Fotion, N.N., Gandal, M.J., et al. (2021). Identification of neural oscillations and epileptiform changes in human brain organoids. Nat. Neurosci. 24, 1488–1500. 10.1038/s41593-021-00906-5.

20. Sharf, T., van der Molen, T., Glasauer, S.M.K., Guzman, E., Buccino, A.P., Luna, G., Cheng, Z., Audouard, M., Ranasinghe, K.G., Kudo, K., et al. (2022). Functional neuronal circuitry and oscillatory dynamics in human brain organoids. Nat. Commun. 13, 4403. 10.1038/s41467-022-32115-4.

21. Quist, E., Trovato, F., Avaliani, N., Zetterdahl, O.G., Gonzalez-Ramos, A., Hansen, M.G., Kokaia, M., Canals, I., and Ahlenius, H. (2022). Transcription factor-based direct conversion of human fibroblasts to functional astrocytes. Stem Cell Rep. 17, 1620–1635. 10.1016/j.stemcr.2022.05.015.

22. Birey, F., Andersen, J., Makinson, C.D., Islam, S., Wei, W., Huber, N., Fan, H.C., Metzler, K.R.C., Panagiotakos, G., Thom, N., et al. (2017). Assembly of functionally integrated human forebrain spheroids. Nature 545, 54–59. 10.1038/nature22330.

23. Schellino, R., Besusso, D., Parolisi, R., Gómez-González, G.B., Dallere, S., Scaramuzza, L., Ribodino, M., Campus, I., Conforti, P., Parmar, M., et al. (2023). hESC-derived striatal progenitors grafted into a Huntington’s disease rat model support long-term functional motor recovery by differentiating, self-organizing and connecting into the lesioned striatum. Stem Cell Res. Ther. 14, 1–21. 10.1186/s13287-023-03422-4.

24. Adler, A.F., Cardoso, T., Nolbrant, S., Mattsson, B., Hoban, D.B., Jarl, U., Wahlestedt, J.N., Grealish, S., Björklund, A., and Parmar, M. (2019). hESC-Derived Dopaminergic Transplants Integrate into Basal Ganglia Circuitry in a Preclinical Model of Parkinson’s Disease. Cell Rep. 28, 3462–3473.e5. 10.1016/j.celrep.2019.08.058.

25. Staresina, B.P., Niediek, J., Borger, V., Surges, R., and Mormann, F. (2023). How coupled slow oscillations, spindles and ripples coordinate neuronal processing and communication during human sleep. Nat. Neurosci. 26, 1429–1437. 10.1038/s41593-023-01381-w.

26. Sanchez-Vives, M.V., and McCormick, D.A. (2000). Cellular and network mechanisms of rhythmic recurrent activity in neocortex. Nat. Neurosci. 3, 1027–1034. 10.1038/79848.

27. Beltramo, R., D’Urso, G., Dal Maschio, M., Farisello, P., Bovetti, S., Clovis, Y., Lassi, G., Tucci, V., De Pietri Tonelli, D., and Fellin, T. (2013). Layer-specific excitatory circuits differentially control recurrent network dynamics in the neocortex. Nat. Neurosci. 16, 227–234. 10.1038/nn.3306.

28. Lőrincz, M.L., Gunner, D., Bao, Y., Connelly, W.M., Isaac, J.T.R., Hughes, S.W., and Crunelli, V. (2015). A Distinct Class of Slow ([0.2–2 Hz) Intrinsically Bursting Layer 5 Pyramidal Neurons Determines UP/DOWN State Dynamics in the Neocortex. J. Neurosci. 35, 5442–5458. 10.1523/JNEUROSCI.3603-14.2015.

29. Watson, B.O., Levenstein, D., Greene, J.P., Gelinas, J.N., and Buzsáki, G. (2016). Network Homeostasis and State Dynamics of Neocortical Sleep. Neuron 90, 839–852. 10.1016/j.neuron.2016.03.036.

30. Moore, A.R., Zhou, W.-L., Sirois, C.L., Belinsky, G.S., Zecevic, N., and Antic, S.D. (2014). Connexin hemichannels contribute to spontaneous electrical activity in the human fetal cortex. Proc. Natl. Acad. Sci. 111. 10.1073/pnas.1405253111.

31. Luhmann, H.J., Sinning, A., Yang, J.-W., Reyes-Puerta, V., Stüttgen, M.C., Kirischuk, S., and Kilb, W. (2016). Spontaneous Neuronal Activity in Developing Neocortical Networks: From Single Cells to Large-Scale Interactions. Front. Neural Circuits 10. 10.3389/fncir.2016.00040.

32. Shaltouki, A., Peng, J., Liu, Q., Rao, M.S., and Zeng, X. (2013). Efficient Generation of Astrocytes from Human Pluripotent Stem Cells in Defined Conditions. Stem Cells 31, 941–952. 10.1002/stem.1334.

33. Wester, J.C., and Contreras, D. (2013). Differential Modulation of Spontaneous and Evoked Thalamocortical Network Activity by Acetylcholine Level In Vitro. J. Neurosci. 33, 17951–17966. 10.1523/JNEUROSCI.1644-13.2013.

34. Fellin, T., Halassa, M.M., Terunuma, M., Succol, F., Takano, H., Frank, M., Moss, S.J., and Haydon, P.G. (2009). Endogenous nonneuronal modulators of synaptic transmission control cortical slow oscillations in vivo. Proc. Natl. Acad. Sci. 106, 15037–15042. 10.1073/pnas.0906419106.

35. Poskanzer, K.E., and Yuste, R. (2011). Astrocytic regulation of cortical UP states. Proc. Natl. Acad. Sci. 108, 18453–18458. 10.1073/pnas.1112378108.

36. Brancaccio, M., Patton, A.P., Chesham, J.E., Maywood, E.S., and Hastings, M.H. (2017). Astrocytes Control Circadian Timekeeping in the Suprachiasmatic Nucleus via Glutamatergic Signaling. Neuron 93, 1420–1435.e5. 10.1016/j.neuron.2017.02.030.

37. Watanabe, A., Guo, C., and Sjöström, P.J. (2023). The developmental profile of visual cortex astrocytes. iScience 26, 106828. 10.1016/j.isci.2023.106828.

38. Szabó, Z., Héja, L., Szalay, G., Kékesi, O., Füredi, A., Szebényi, K., Dobolyi, Á., Orbán, T.I., Kolacsek, O., Tompa, T., et al. (2017). Extensive astrocyte synchronization advances neuronal coupling in slow wave activity in vivo. Sci. Rep. 7, 6018. 10.1038/s41598-017-06073-7.

39. de Ceglia, R., Ledonne, A., Litvin, D.G., Lind, B.L., Carriero, G., Latagliata, E.C., Bindocci, E., Di Castro, M.A., Savtchouk, I., Vitali, I., et al. (2023). Specialized astrocytes mediate glutamatergic gliotransmission in the CNS. Nature 622, 120–129. 10.1038/s41586-023-06502-w.

40. Vaidyanathan, T.V., Collard, M., Yokoyama, S., Reitman, M.E., and Poskanzer, K.E. (2021). Cortical astrocytes independently regulate sleep depth and duration via separate GPCR pathways. eLife 10, e63329. 10.7554/eLife.63329.

41. Crunelli, V., and Hughes, S.W. (2010). The slow (<1 Hz) rhythm of non-REM sleep: a dialogue between three cardinal oscillators. Nat. Neurosci. 13, 9–17. 10.1038/nn.2445.

42. Timofeev, I., Grenier, F., and Steriade, M. (2000). Impact of intrinsic properties and synaptic factors on the activity of neocortical networks in vivo. J. Physiol.-Paris 94, 343–355. 10.1016/S0928-4257(00)01097-4.

43. Lörincz, M.L., Crunelli, V., and Hughes, S.W. (2008). Cellular Dynamics of Cholinergically Induced α (8–13 Hz) Rhythms in Sensory Thalamic Nuclei *In Vitro*. J. Neurosci. 28, 660–671. 10.1523/JNEUROSCI.4468-07.2008.

44. Buzsáki, G. (2001). Hippocampal GABAergic Interneurons: A Physiological Perspective. Neurochem. Res. 26, 899–905. 10.1023/A:1012324231897.

